# In vivo transition in chromatin accessibility during differentiation of deep-layer excitatory neurons in the neocortex

**DOI:** 10.1101/2024.08.26.609258

**Authors:** Seishin Sakai, Yurie Maeda, Keita Kawaji, Yutaka Suzuki, Yukiko Gotoh, Yusuke Kishi

## Abstract

During the differentiation process of neurons, their gene transcription pattern changes according to both intrinsic and extrinsic stimuli-induced programs. Chromatin regulation at regulatory elements is involved in this precise control of gene expression. However, developmental changes in chromatin accessibility of the cortical neurons in vivo are less understood, partly because there is no convenient method to genetically label neurons of a specific lineage. Here, we establish a method for labeling the differentiating neurons of specific birthdates. Using this method, we traced the four-day differentiation process of in vivo deep-layer excitatory neurons in mouse embryonic cortex and examined the changes in the genome-wide transcription pattern and chromatin accessibility with RNA-seq and DNase-seq, respectively. The genomic regions of genes associated with mature neuron functions, such as deep layer–specific genes and genes responsive to external stimuli, became open even in the embryonic stage. Moreover, genes with bivalent marks in neural precursor/stem cells (NPCs) became open. Together, our data demonstrate the importance of chromatin regulation in vivo differentiating neurons during the embryonic stage to follow activation of neuronal genes in their maturation process.

## Introduction

During cortical development, distinct subtypes of neurons are sequentially generated from common neural precursor/stem cells (NPCs) residing in the ventricular zone (VZ) according to their birthdate (Molyneaux et al., 2007). After neuronal fate commitment, they migrate to the pial surface and constitute a six-layered structure, that are composed of different subtypes of neurons, with an inside–out pattern. After completing migration, they mature and acquire neuronal functions, such as neuronal subtype- and layer-specific features and transcription profiles. For example, layer 6 neurons project their axons into the thalamus and are also implicated as pioneer neurons in the callosal projection (Carlos and O’Leary, 1992; O’Leary and Koester, 1993). Considering that the neuronal subtypes produced by NPCs change day-to-day in the mouse neocortex, it is necessary to label, isolate, and analyze in vivo the specific lineage of neurons to precisely understand the neuronal differentiation process. Transcriptomic changes during neuronal differentiation of each subtype were revealed by a combination of the original labeling system, Flash-tag, and single-cell RNA-seq (scRNA-seq) (Telley et al., 2019, 2016).

Among several layers of transcriptional regulation, chromatin accessibility has a role in establishing a preparatory state for transcription factors. Transcription factors can regulate genes in accessible and open chromatin, but have difficulty in inaccessible and closed chromatin (Boyle et al., 2008a; Crawford et al., 2006). Several studies have shown that chromatin opening during the developmental process serves as the basis for following activation in mature cell types. For instance, in intestinal progenitors, regulatory elements determined by dimethylation of histone H3 at lysine 4 (H3K4me2) are already opened (Kim et al., 2014). In the hematopoietic system, binding sites of master transcription factors, such as Foxp3 in regulatory T cells or PU.1 in granulocytes, also became accessible before they were expressed (Luyten et al., 2014; Samstein et al., 2012). Recent single-cell multiome analysis, acquiring both gene expression and chromatin accessibility data from the same single cells, also revealed the preceding opening of chromatin-related lineage specifications during the differentiation process in hair follicles (Ma et al., 2020).

In the central nervous system, several studies have shown the role of chromatin accessibility in vivo in the developing brain. One pioneering study using DNase-seq, a method for profiling chromatin-accessible genomic regions by a DNaseI nuclease, revealed that the chromatin opening mediated by Zic transcription factors contributes to transcriptional activation during in vivo differentiation of cerebellar granule neurons (Frank et al., 2015). A recent single-cell assay for transposase-accessible chromatin with high-throughput sequencing (ATAC-seq) analyses using mouse and human developing brain also indicates the role of chromatin openness at the regulatory element of neuronal genes on the following gene transcription during neuronal differentiation (Bella et al., 2021; Noack et al., 2022; Trevino et al., 2021; Ziffra et al., 2021). However, these studies analyzed all cell types of developing brains at only one or a small number of stages and assumed the lineage relationship; thus, changes in chromatin accessibility during the differentiation of a single lineage of neurons remain unknown.

Here, we develop a novel method for genetically labeling a specific lineage of cortical neurons by in utero electroporation using wild-type mice and trace embryonic differentiation of in vivo deep-layer neurons that were labeled. Taking advantage of our method, the changes in transcriptome and chromatin accessibility were revealed by RNA-seq and DNase-seq simultaneously during their differentiation process. In our analysis, we found the genes with the promoter regions that became open during neuronal differentiation in the embryonic stage were enriched with deep layer–specific genes and responsive genes for brain-derived neurotrophic factor (BDNF) stimulation, indicating that the regulation of the chromatin accessibility might contribute to acquiring a preparatory state for the induction of genes related to mature neuronal functions. Furthermore, we also found that chromatin opening during embryonic neuronal differentiation took place mainly in bivalent genes in NPCs that had both H3K4me3 and H3K27me3 histone modifications.

## Results

### In vivo tracing of neuronal differentiation of specific birthdate in neocortex by genetical labeling with in utero electroporation

To understand neuronal differentiation in the neocortex, we first attempted to genetically label a specific lineage of neurons determined by their specific birthdate since neocortical NPCs sequentially produce different types of neurons in the embryonic stage. We introduced two plasmids, pAAV-DIO-GFP and pNeuroD1-ERT2CreERT2, into NPCs by in utero electroporation. pAAV-DIO-GFP expresses GFP after the activation of Cre recombinase since the GFP cassette is inversely inserted between two types of loxP sequence (Schnütgen et al., 2003). Since neurons transiently express *NeuroD1* in their immature state after finishing cell devision (Miyoshi and Fishell, 2012), pNeuroD1-ERT2CreERT2 expresses tamoxifen-activating Cre recombinase only in NeuroD1-positive immature neurons (Guerrier et al., 2009). Therefore, we can permanently label a specific lineage of immature neurons with the same birthdate when tamoxifen is injected (Figure 1a). To confirm this method, we introduced these two plasmids together with mCherry-expressing plasmid under the control of the ubiquitous promoter CAG, as a transfection control at embryonic day 12.0 (E12.0) neocortex and injected tamoxifen at E13.0 (Figure 1b). Immunostaining of embryos at E16.0 showed that GFP-labeled cells were mainly located in the deep layer labeled with Ctip2, a marker for deep-layer neurons (Arlotta et al., 2005). By contrast, mCherry-positive cells are distributed not only in the cortical plate (CP) but also in the intermediate zone (IMZ) and even in the ventricular and subventricular zone (VZ/SVZ). This result suggests that neurons produced at later stage, at least four days after transfection, were also labeled with a plasmid of ubiquitous promoter. In contrast, our tracing method specifically labeled Ctip2-positive and deep-layer neurons produced before the tamoxifen injection.

**Figure 1.**
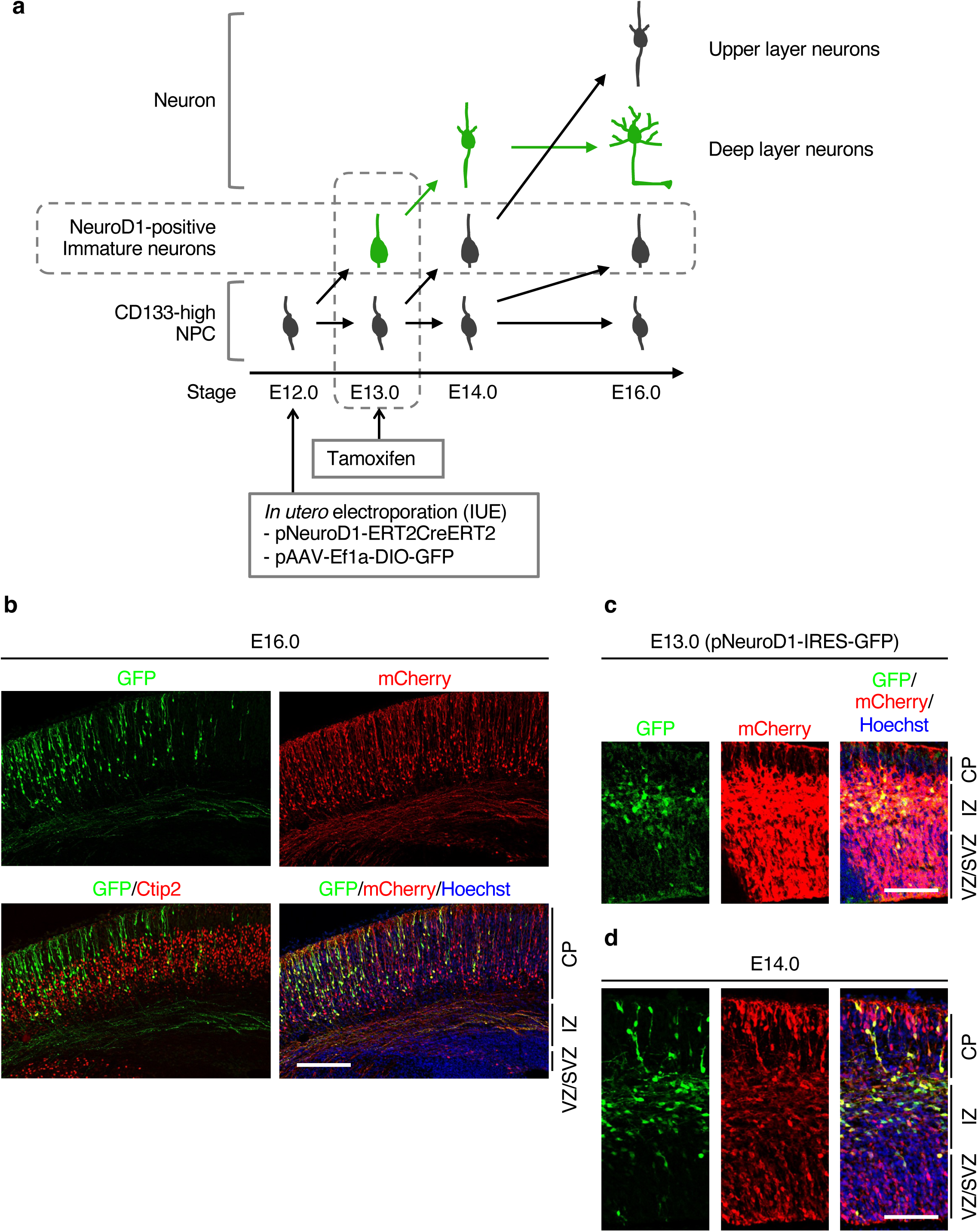
Genetic labeling of deep-layer neurons by in utero electroporation. **(a)** An experimental scheme is used to label deep-layer neurons and trace their differentiation process. (**b–d**) pNeuroD1-ERT2CreERT2, pAAV-Ef1a-DIO-GFP, and pCAG-mCherry plasmids (b, d) or pNeuroD1-IRES-GFP and pCAG-mCherry plasmids (c) were injected into the lateral ventricle of E12.0 embryos and electroporated. The pregnant mice were injected with tamoxifen at E13.0 (b, d). The brains were dissected out from the uterus at indicated stages and subjected to immunohistofluorescence staining with anti-GFP, anti-RFP (mCherry), and anti-Ctip2 (only b) antibodies. Nuclei were counterstained with Hoechst 33342. Representative images are indicated. Scale bars, 400 mm (b) and 200 mm (c, d). VZ/SVZ, ventricular/subventricular zone. IZ, intermediate zone. CP, cortical plate.

We next confirmed the differentiation status of the labeled neurons at E13.0 and E14.0. Since tamoxifen injection was not yet performed at E13.0, we visualized neurons to be labeled by transfection of pNeuroD1-IRES-GFP (Figure 1c) and found that GFP-positive cells were mainly located in IMZ. At E14.0, one day after tamoxifen injection, GFP-positive neurons were primarily retained in IMZ, but some of them moved to CP (Figure 1d). Therefore, this labeling method allows us to trace the differentiation process of deep-layer neurons with a time resolution of one day.

To examine their gene expression patterns, we next isolated these cells by fluorescent-activated cell sorting (FACS) at E14.0 and E16.0 together with CD133-high NPCs at E12.0 as the originating progenitor cells for labeled cells. pNeuroD1-IRES-GFP positive cells were sorted at E13.0, as neurons that were labeled by tamoxifen injection in our system (Figure 2a). Reverse transcription and quantitative polymerase chain reaction (RT-qPCR) analysis confirmed that the expression level of *Nestin*, an NPC marker, reduced, whereas those of *Tubb3*, *Gabrb2*, neuronal markers, increased in pNeuroD1-ERT2CreERT2–labeled cells at E16.0 compared to CD133-high cells at E12.0 or pNeuroD1-IRES-GFP–positive cells at E13.0 (Figure 2b). Consistent with the pNeuroD1 activity, endogenous *NeuroD1* was highly expressed in pNeuroD1-IRES-GFP–positive cells at E13.0. These data again suggest that the embryonic differentiation process of deep-layer excitatory neurons was successfully traced in our labeling system. Then, CD133-high cells at E12.0, pNeuroD1-IRES-GFP–positive cells at E13.0, and pNeuroD1-ERT2CreERT2–labeled cells at E14.0 and E16.0 are referred to as E12.0 NPCs, E13.0 neurons, and E14.0 and E16.0 neurons, respectively.

**Figure 2.**
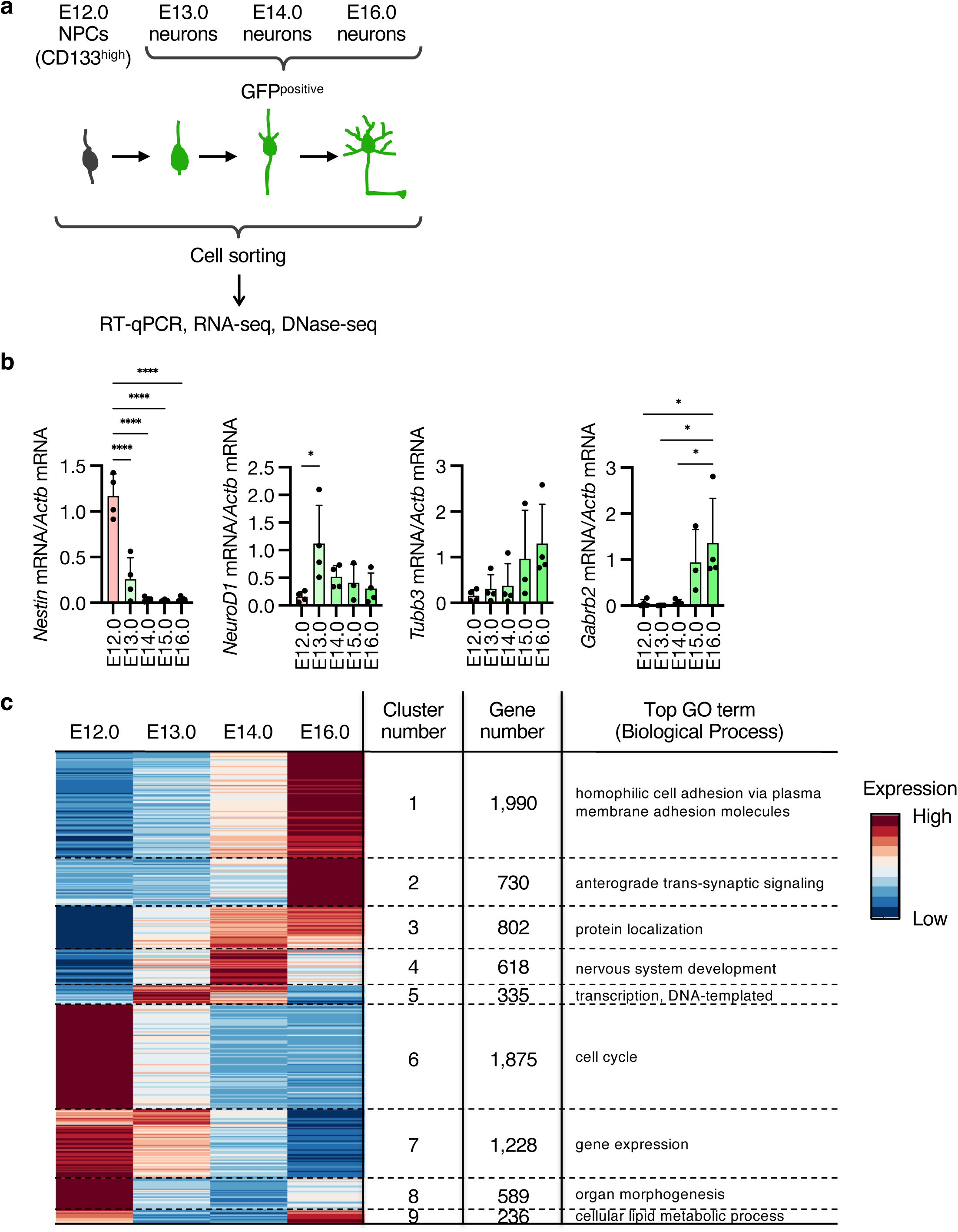
Transcriptomic changes during 4-day differentiation of deep-layer neurons. **(a)** Experimental scheme to isolate E12.0 NPCs and GFP-labeled neurons at E13.0, E14.0, and E16. (**b**) RT-qPCR analysis of isolated NPCs and neurons. Relative abundance of indicated mRNA (normalized by the amount of Actb mRNA) was shown. Data are means + SD from three (E15.0) or four (E12.0, E13.0, E14.0, and E16.0) independent experiments. (**c**) RNA-seq analysis of isolated E12.0 NPCs and E13.0, E14.0, and E16.0 neurons. Expression levels of DEGs between four cell types were shown in the heatmap. The DEGs were categorized into seven clusters, and the gene numbers and the top GO term in the Biological Process for each gene cluster are shown.

Next, we performed an RNA-seq analysis with E12.0 NPCs and E13.0, E14.0, and E16.0 neurons. Developmentally regulated genes during differentiation from E12.0 NPCs to E16.0 neurons were determined and classified into 9 clusters by maSigPro software (FDR < 0.05) (Conesa et al., 2006). The transcription levels of more than half of the expressed protein-coding genes (8,403 genes out of 14,286 expressed genes) significantly changed (Figure 2c), indicating the global changes of transcription patterns in four days of differentiation from NPCs to embryonic neurons. In upregulated genes, genes related to neuronal functions, such as synaptic transmission and protein transport, were specifically enriched (Cluster 2) (Figure 2c). On the other hand, genes related to the cell cycle were enriched in downregulated genes (Cluster 6), indicating that the cell cycle exits at the onset of neuronal differentiation. Notably, genes related to the lipid biosynthetic process were specifically enriched in the cluster of temporally suppressed genes (Cluster 9) (Figure 2c). This suggests that the synthesis of lipids, essential components for membrane, is important for the future maturation process of neurons, such as elaborating neurites, in addition to supplying lipids for the cell cycle–associated increase of plasma membrane. Overall, these results suggest the dynamic change in transcription states during only four days of differentiation from NPCs in the embryonic stage in vivo.

### Changes of DNase-hypersensitive sites (DHSs) during in vivo differentiation of deep-layer neurons

Since the regulation of chromatin accessibility at gene regulatory sites is known to be important for the regulation of the transcription states of target genes, we next performed DNase-seq using E12.0 NPCs and E13.0, E14.0, and E16.0 neurons (Figure 3a). Isolated nuclei were digested by DNaseI, and DNaseI sensitive fragments, shorter than 1 kbp, and were collected by gel extraction after agarose gel electrophoresis at each stage. The sequence of each collected DNA fragment was determined by high-throughput sequence, and DHSs were determined by f-seq software (Boyle et al., 2008b) (Figure 2b). In our analysis, approximately 20,000–30,000 DHSs were determined at each stage, and E16.0 neurons had the most DHSs compared to other stages. The total width of DHSs was higher in E12.0 NPCs and E16.0 neurons compared to E13.0 and E14.0 neurons (Figure 3b), suggesting wider DHSs in E12.0 NPCs. To confirm whether our DNase-seq appropriately identified open chromatin regions, we examined the distribution of DHSs in the genome and the relationship between the openness of gene loci and their expression levels determined by RNA-seq. Consistent with previous reports (Boyle et al., 2008a), DHSs determined by our DNase-seq were enriched in promoter regions, defined by −1 kb to +0.5 kb of transcription start sites (TSS), and exon regions compared to their proportion in the whole genome (Figure 3c). We then determined genes with at least one DHS in their promoter as DHS-positive genes and those without any DHSs in their promoter as DHS-negative genes. As expected, the transcription levels of DHS-positive genes were higher than those of DHS-negative genes at every differentiation stage (Figure 3d). These results suggest that our DNase-seq determined accessible chromatin regions that are enriched in gene coding regions, especially in promoter regions and in active gene loci, as previously reported (Boyle et al., 2008a).

**Figure 3.**
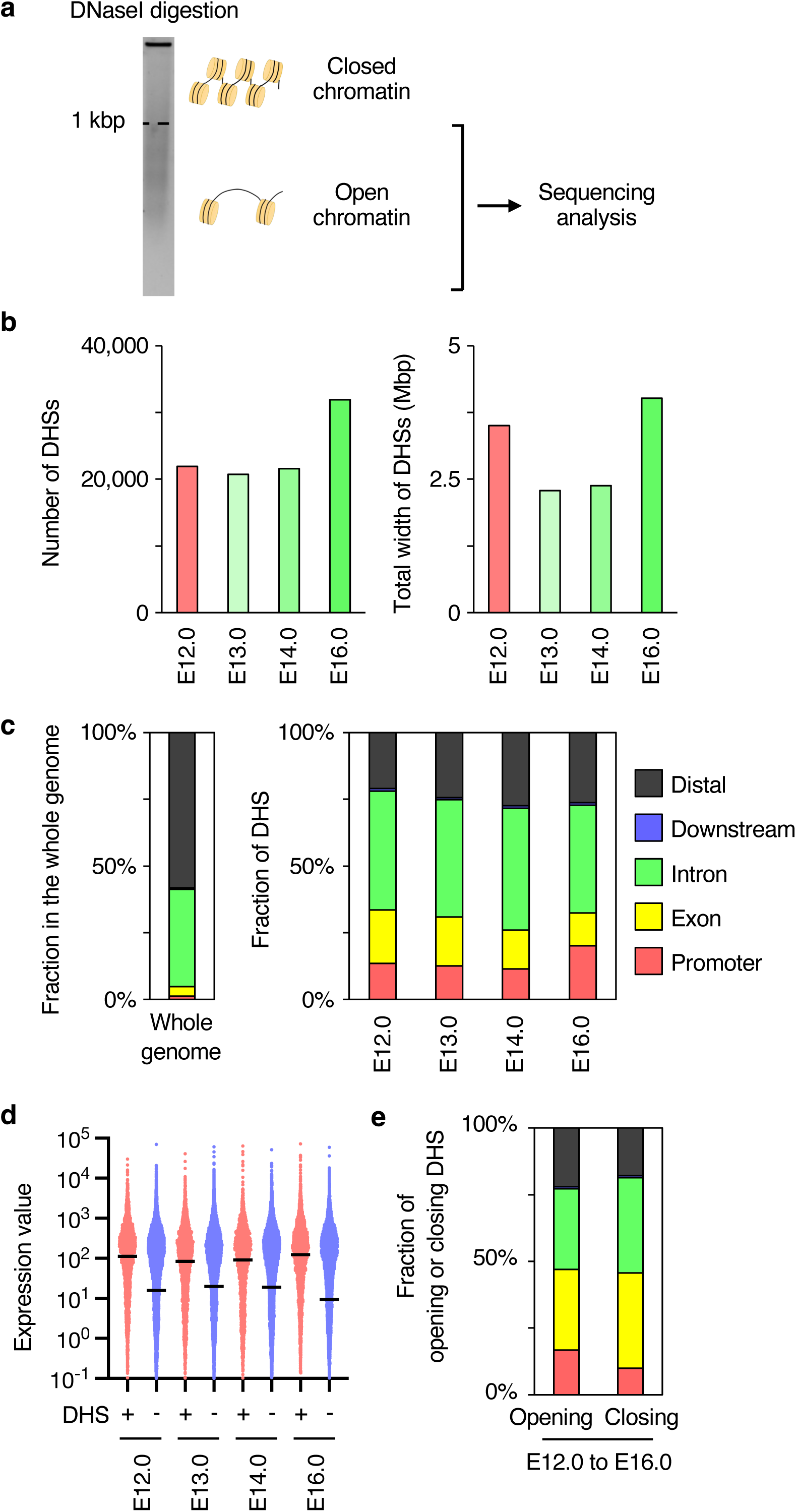
Changes in chromatin accessibility during 4-day differentiation of deep-layer neurons. **(a)** Experimental scheme of DNase-seq analysis. (**b**) The numbers (left panel) and the total width (right panel) of DHSs are identified at each time point. (**c**) The fractions of genomic features in the whole mouse genome (left panel) and DHSs at each time point. (**d**) Expression levels of genes with or without DHSs at each time point. (**e**) The fractions of genomic features in DHSs of opening or closing from E12.0 NPCs to E16.0 neurons.

We next examined the transition of DHSs during in vivo differentiation of deep-layer neurons and found that opening chromatin regions from E12.0 NPCs to E16.0 neurons were specifically enriched in promoter regions compared to closing regions (Figure 3e). This is consistent with the previous reports that neuron-specific accessible regions were especially enriched in promoters (Torre-Ubieta et al., 2018).

Therefore, we next focused on the genes with opening or closing promoters during differentiation. We defined the genes whose DHSs newly appeared in their promoter during differentiation as opening genes and the genes with all the DHSs absent from their promoter during differentiation as closing genes (Figure 4a, b). The transcription levels of opening genes were significantly upregulated during differentiation from E12.0 NPCs to E16.0 neurons, suggesting that the chromatin regulation at the promoter correlates with their changes in transcription levels (Figure 4a). GO analysis showed the enrichment of genes for neuronal functions in the opening gene set (Figure 4c), suggesting that the chromatin opening at promoter regions during differentiation is important for neuronal differentiation. By contrast, the expression levels of closing genes did not significantly change (Figure 4b). These closing genes were enriched with the genes for the metabolic process as revealed by GO analysis (Figure 4d). Given the amount of metabolites with highly unsaturated structures decreases during stem cell differentiation (Yanes et al., 2010), the closing of the promoter regions might contribute to the different usage of metabolic processes between NPCs and neurons.

**Figure 4.**
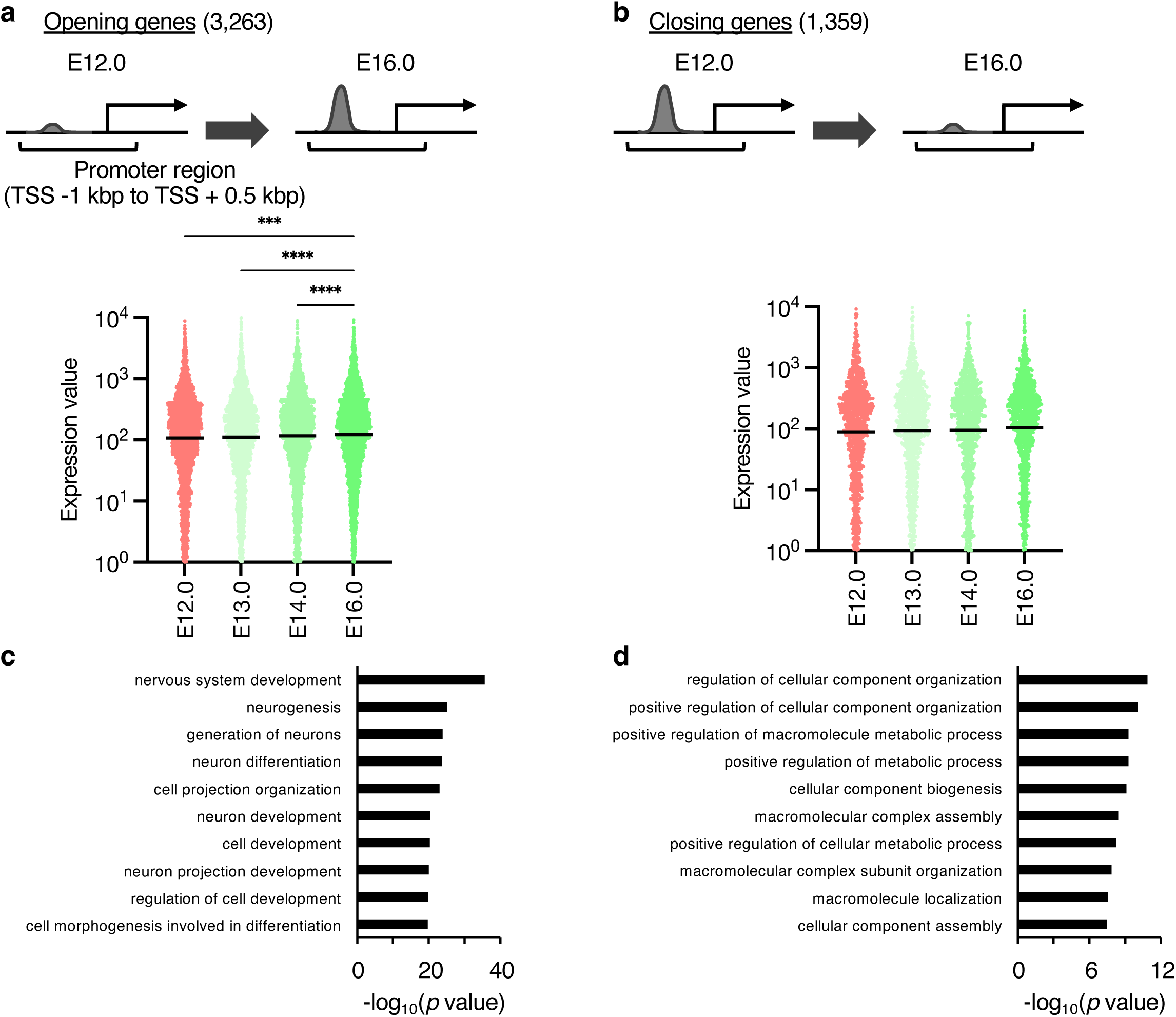
Chromatin opening at promoter regions of neuronal genes during 4-day differentiation of deep-layer neurons. (**a, b**) The expression levels of opening (a) or closing (b) genes at their promoter regions from E12.0 to E16.0. *p* values were determined by the Friedman test, followed by Dunn’s multiple comparison test. \*\*\**p* < 0.001, \*\*\*\**p* < 0.0001. (**c, d**) The top ten GO terms and their *p*-values of opening (c) or closing (d) genes at their promoter regions from E12.0 to E16.0.

Taken together, we successfully acquired the data for the transition of chromatin accessibility during the initial phase of neuronal differentiation in vivo. Using these data, we examined the role of the changes in chromatin accessibility on the transcription of neuronal genes in mature neurons (Figure 5) and the molecular mechanisms for the chromatin opening during neuronal differentiation (Figure 6).

**Figure 5.**
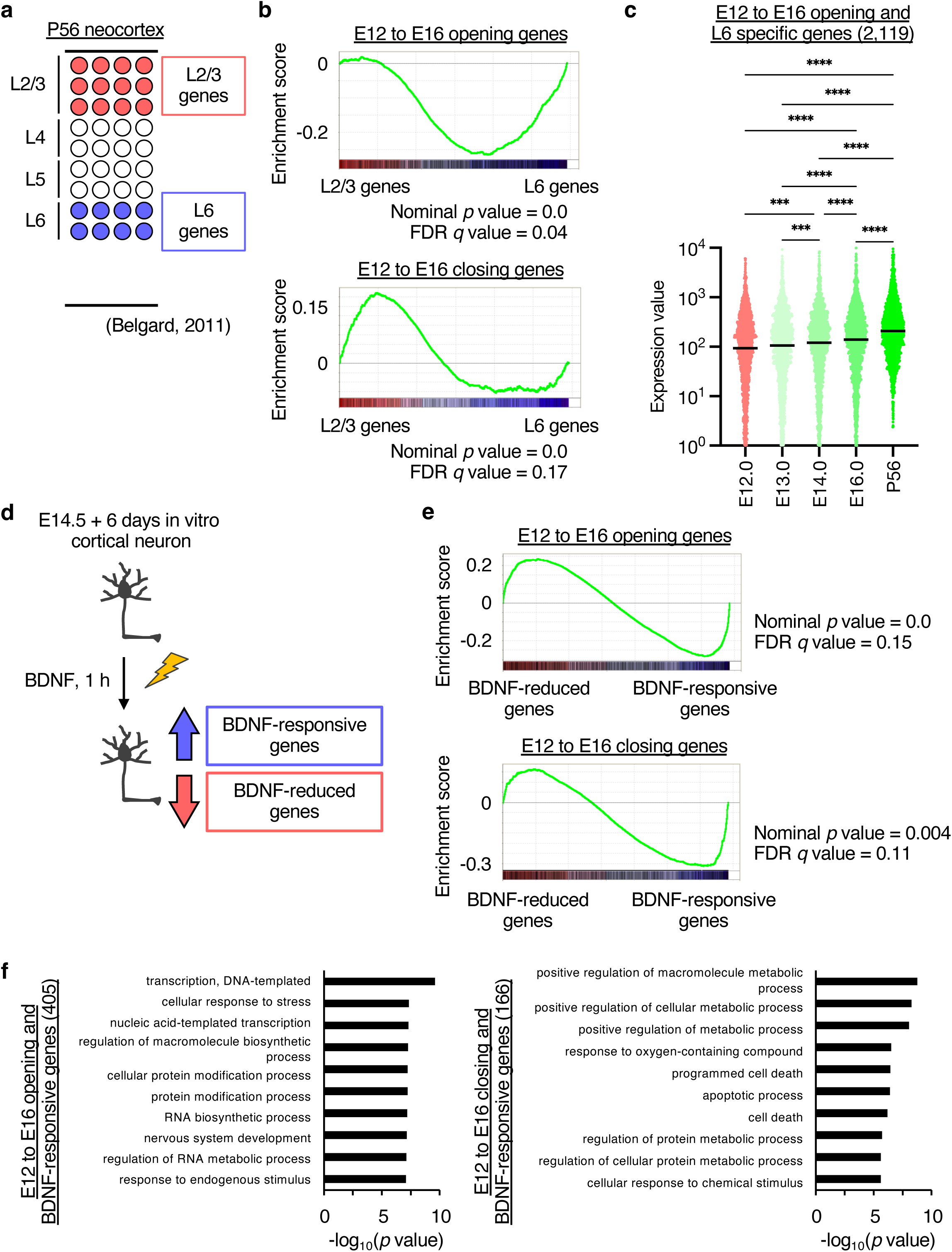
The association of opening genes from E12.0 NPCs to E16.0 neurons and deep-layer specific or BDNF-responsive genes. **(a)** RNA-seq analysis of layer 2/3 (upper layer) and layer 6 (deep layer) at P56 (Belgard et al., 2011). (**b**) GSEA of opening (upper panel) or closing (lower panel) genes from E12.0 NPCs to E16.0 neurons in RNA-seq of layers 2/3 and 6. (**c**) Expression levels of E12.0 NPCs to E16.0 opening and layer 6–specific genes during differentiation from E12.0 NPCs to P56 neurons. *p*-values were determined by the Friedman test, followed by Dunn’s multiple comparison test. \*\*\**p* < 0.001, \*\*\*\**p* < 0.0001. (**d**) Microarray analysis of in vitro primary culture neurons with or without BDNF stimulation for 1 h. (**e**) GSEA of opening (upper panel) or closing (lower panel) genes from E12.0 NPCs to E16.0 neurons in vitro primary culture neurons with or without BDNF stimulation. (**f**) GO analysis of E12.0 NPCs to E16.0 opening (left panel) or closing (right panel) and BDNF-responsive genes.

**Figure 6.**
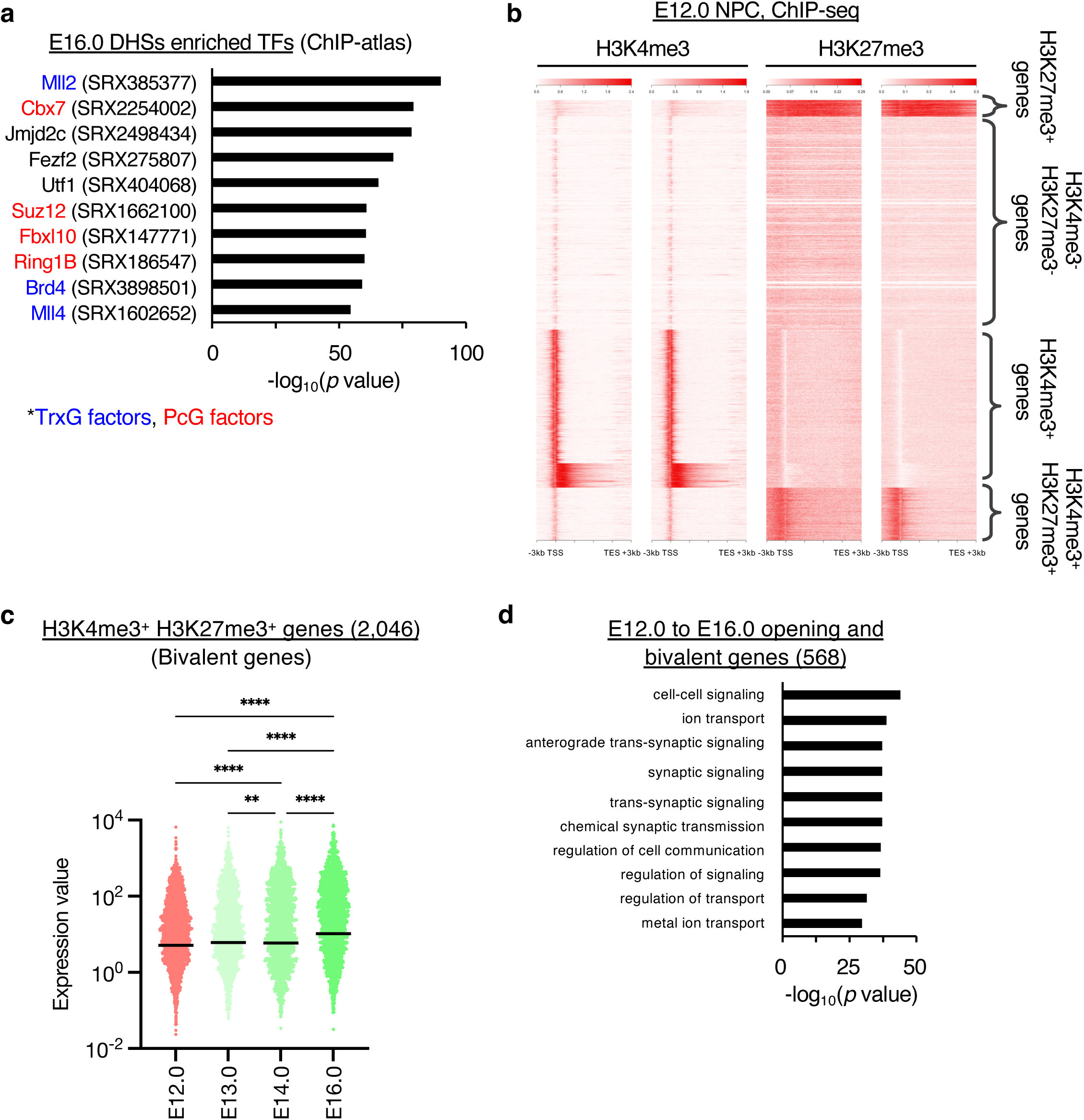
The association between chromatin opening during neuronal differentiation and bivalent state in NPCs. **(a)** Transcription factors enriched in DHSs of E16.0 neurons by ChIP-atlas analysis. (**b**) E12.0 NPCs were subjected to ChIP-seq analysis with H3K4me3 and H3K27me3 antibodies. All genes were categorized into five clusters on the basis of the pattern of H3K4me3 and H3K27me3 around gene regions. (**c**) Expression levels of H3K4me3-positive and H3K27me3-positive bivalent genes in E12.0 NPCs during 4-day differentiation. *p*-values were determined by the Friedman test, followed by Dunn’s multiple comparison test. \**p* < 0.05, \*\**p* < 0.001 \*\*\*\**p* < 0.0001. (**d**) GO analysis of E12.0 NPCs to E16.0 opening and bivalent genes in E12.0 NPCs.

### Acquiring chromatin accessibility during embryonic differentiation for the preparatory state of neuronal gene loci

Chromatin accessibility contributes not only to the transcriptional activity at the timing but also to the potential of the future induction of the genes. Indeed, previous reports found an increase in chromatin accessibility in the gene loci that would be activated or bound by transcription factors during the developmental process of neural and non-neural systems (Frank et al., 2015; Kim et al., 2014; Luyten et al., 2014; Samstein et al., 2012). In order to examine whether the regulation of chromatin accessibility plays any role in the induction of genes for deep-layer excitatory neurons, we examined the chromatin state of neuronal gene sets that are mostly activated in the postnatal and adult stages. We particularly focused on the layer-specific genes and responsive genes for external stimuli.

Cortical neurons have layer-specific functions, such as their specific morphology and target brain regions. Deep-layer neurons, including layer 5 and 6 neurons, the main layers that we were analyzing in this study, also have specific functions (Arlotta et al., 2005; Carlos and O’Leary, 1992; Molyneaux et al., 2007; O’Leary and Koester, 1993). To examine the chromatin accessibility of layer-specific genes, we utilized previously reported RNA-seq analysis (Figure 5a) (Belgard et al., 2011) and investigated the enrichment of opening and closing genes from E12.0 NPCs to E16.0 neurons by gene set enrichment analysis (GSEA) (Mootha et al., 2003; Subramanian et al., 2005). Opening genes were significantly enriched in layer 6–specific genes, whereas closing genes were enriched in layer 2/3–specific genes (Figure 5b). Given that isolated neurons mostly differentiate into deep-layer excitatory neurons (Figure 1b), these data suggest that the preparatory state for deep layer–specific gene expression patterns has already been established during the embryonic differentiation process. In addition, the expression levels of deep-layer specific genes that became open during differentiation from E12.0 NPCs to E16.0 neurons slightly increased from E12.0 NPCs to E16.0 neurons, whereas they were obviously upregulated from E16.0 neurons to layer 6 neurons 56 days post-natal (P56) (Figure 5c) (Belgard et al., 2011). This further supports the idea that preparatory state for deep layer–specific gene expression patterns has already been established in the embryonic stage.

Neuronal activity induced by external stimuli, especially in the postnatal stage, is important for the functional maturation of neurons through regulating neurite extension and synapse formation (West and Greenberg, 2011). In this process, the expression of responsive genes for external stimuli, such as neurotrophic factors, plays a crucial role. Therefore, we next focused on responsive genes for external stimuli. To define these gene sets, we performed a microarray analysis of BDNF-stimulated in vitro primary neurons. In vitro neurons cultured for six days from E15 cortex were stimulated with BDNF for 1 h. Using the results of the microarray, GSEA analysis for opening genes from E12.0 NPCs to E16.0 neurons showed significant enrichment in upregulated genes by BDNF stimulation (Figure 5e). Furthermore, genes showing both opening during differentiation and upregulation by BDNF-stimulation specifically included genes related to neuronal development determined by GO analysis (Figure 5f). Closing genes from E12.0 NPCs to E16.0 neurons were also enriched in upregulated genes by BDNF stimulation (Figure 5e). However, these genes included genes related to apoptosis, suggesting excitotoxicity by BDNF stimulation in an in vitro culture (Figure 5f). Overall, these results support the notion that chromatin opening during embryonic differentiation confers the permissiveness for the induction of genes associated with neuronal functions, such as layer-specific genes and responsive genes to neurotrophic factors.

### Opening of bivalent gene loci in NPCs during neuronal differentiation

Another key question is what regulates the transition in chromatin accessibility during neuronal differentiation. To address this, we investigated the transcription factors binding specifically to DHSs determined in E16.0 neurons rather than those in E12.0 NPCs, using ChIP-atlas, a database of the binding sites of various transcription factors and chromatin regulators determined using public ChIP-seq analysis (Oki et al., 2018). We found binding sites of several factors were enriched in DHSs of E16.0 neurons (Figure 6a). Among these, we focused on the chromatin regulators of both Trithorax group proteins (TrxG), such as Mll family proteins and Brd4, and Polycomb group proteins (PcG), such as Cbx7, Suz12, Fbxl10, and Ring1B (Schuettengruber et al., 2017). In embryonic stem (ES) cells, genes that harbor both H3K4me3 modified by Mll in TrxG and H3K27me3 modified by Ezh2 in PcG are called bivalent genes and known to be poised for activation or inactivation during subsequent differentiation process to specific lineage (Azuara et al., 2006; Bernstein et al., 2006; Zhao et al., 2007).

Therefore, to examine the chromatin regulation of bivalent genes in NPCs, we determined the bivalent gene set in E12.0 NPCs by ChIP-seq analysis for H3K4me3 and H3K27me3 (Figure 6b). Unbiased clustering analysis with patterns of these histone modifications around gene body regions determined bivalent genes with both H3K4me3 and H3K27me3 around TSS regions. Consistent with the results of the ChIP-atlas (Figure 6a), a portion of bivalent NPCs became open during neuronal differentiation (568 genes out of 2,046 bivalent genes). Importantly, among these four gene sets, the expression levels of bivalent genes increased during neuronal differentiation (Figure 6c). GO analysis also revealed the enrichment of genes related to mature neuronal functions, such as synaptic signaling and ion transport, in bivalent genes in NPCs (Figure 6d). Overall, these results suggest that the opening of bivalent genes in E12.0 NPCs contributes to the upregulation of neuronal genes during embryonic neuronal differentiation.

## Discussion

The competence of neurons for their functions, such as neuronal subtype-specific functions and responsiveness to external stimuli, is established during the neuronal differentiation process from NPCs. Although many studies have examined changes by biochemical analysis with in vitro primary culture of neurons, the neuronal differentiation process in vitro is very different than that in vivo, that involves neuronal migration and drastic morphological changes. Since brains, even in the embryonic stage, consist of many cell types, including different subtypes of neurons, glial cells, and non-neural cells, it is necessary to label and isolate specific lineages of neurons to analyze the neuronal differentiation process in vivo in the brain. Telley et al. reported the Flash-Tag method to label cortical neurons with the same birthdate by random conjugation of fluorescent dye to intracellular proteins (Telley et al., 2016). Flash-Tag is very useful because of its simplicity; however, the proteins labeled with fluorescence degrade and shorten the tracing duration time. Using Neurog2-CreERT2 transgenic mice, Hirata et al. succeeded in genetically and permanently labeling cortical neurons with the same birthdate (Hirata et al., 2021, 2016; Hou et al., 2019). Our novel methods introducing pNeuroD1-ERT2CreERT2 by in utero electroporation have advantages when using wild-type mice and labeling permanently cortical neurons with the same birthdate.

Leveraging these advantages, we traced the differentiation of neurons produced from NPCs around E12 for 4 days and isolated them for performing RNA-seq and DNase-seq analyses simultaneously. Seventy-five percent of differentially expressed genes (1,742 genes out of 2,318 genes) observed by Telley et al. were also annotated as differentially expressed genes in our analysis, indicating the feasibility of our new labeling method in labeling newborn neurons at similar stages (Telley et al., 2016). In addition to this, our data revealed that the transcription levels of more than half of protein-coding genes significantly changed during neuronal differentiation, indicating the global changes in transcription states during neuronal differentiation in vivo as observed with in vitro primary cultures (Kaur et al., 2014).

Chromatin openness, as examined by DNase-seq, has an important role in gene transcription. In our analysis, genomic regions with chromatin openness during neuronal differentiation were enriched in promoter regions, and the changes at the promoter were correlated with the change in transcription levels of nearby genes, indicating the contribution of chromatin regulation to a change in transcription patterns. This differs from the results of the ATAC-seq of granule neurons in the dentate gyrus with electroconvulsive stimulation showing that the most changes in chromatin openness were observed in the intron and distal regions (Su et al., 2017). Therefore, the regulation of chromatin might differ between developmental and transient changes in gene transcription. In addition, given that the chromatin regulation at the enhancer also contributes to the regulation of the transcription state during development (Heintzman et al., 2009; Song et al., 2011), it is possible that the chromatin regulation at distal regulatory sites is also important for the regulation of gene transcription during neuronal differentiation.

In addition to transcriptional activity at the timing, chromatin opening also contributes to the preparatory state for future activation. Here, our results suggest that the chromatin opening without gene activation in the gene loci is associated with the features of deep-layer excitatory neurons, such as the deep layer–specific genes and BDNF stimulus-dependent genes, during the embryonic differentiation process. Interestingly, closing genes from NPCs to E16 neurons were enriched in upper layer–specific genes. These results suggest that chromatin regulation during the embryonic stage confers layer-specific neuronal functions. In the case of responsive genes for external stimuli, previous reports have shown that electroconvulsive stimulation to granule neurons in the dentate gyrus induces chromatin opening mainly in distal regulatory elements of responsive genes in a c-Fos–dependent manner (Su et al., 2017). Combined with our results, responsive genes may acquire a preparatory state by opening their promoter regions in the embryonic stage and activating their transcription through their enhancer regions.

In order to identify the chromatin regulator during neuronal differentiation, we took advantage of ChIP-atlas database for the binding sites of the transcription factors determined by ChIP-seq experiments (Oki et al., 2018). We found that the binding sites for the histone methyltransferase, such as Mll proteins for H3K4me3 and components of PcG for H3K27me3, involved in bivalent marks determined in ES cells, were specifically enriched in DHSs determined in E16 neurons. Our ChIP-seq analyses for H3K27me3 and H3K4me3 with NPCs showed that bivalent genes in NPCs became active during neuronal differentiation from NPCs to E16 neurons. This result suggests that the opening of bivalent genes contributes to the activation of neuronal genes. Previous studies proposed several mechanisms for the activation of bivalent genes, such as the contribution of H3K27me3 demethylase and chromatin remodeler complex (Dhar et al., 2016; Narayanan et al., 2015; Tang et al., 2020). Our motif analysis for the opening and bivalent regions during differentiation from E12.0 NPCs to E16.0 neurons by HOMER software revealed the enrichment of the DNA sequence, which is similar to the binding motif of the Dmrt3 protein (Supplementary Figure 1a). Previous reports have shown that Dmrt3 and Dmrta2 (also known as Dmrt5) are important for maintaining the undifferentiated state of NPCs by repressing neurogenic genes (Desmaris et al., 2018; Konno et al., 2019, 2012). Given that Dmrt3 represses target genes together with Dmrta2 (Desmaris et al., 2018; Konno et al., 2019), downregulation of Dmrt3 and Dmrta2 might trigger the opening and upregulation of bivalent genes during differentiation from E12.0 NPCs to E16.0 neurons. The expression of Dmrt proteins is regulated downstream of the Wnt and Bmp signaling pathways (Clercq et al., 2016; Desmaris et al., 2018; Konno et al., 2012; Saulnier et al., 2012). Therefore, Dmrt proteins might be important for each NPC to modulate the timing for the activation.

In this study, we have described the transcriptomic and chromatin changes during the differentiation of deep-layer neurons in mouse neocortex by a novel labeling method for a specific lineage of neurons, and the opening of the chromatin state for preparing future gene activation. Our findings provide a basis for future studies to reveal the mechanisms for acquiring neuronal functions during their development.

## Methods

### Ethics statement

All animals were maintained and studied according to protocols approved by the Animal Care and Use Committee of The University of Tokyo (approval numbers: P25-8 and P30-4 in Graduate School of Pharmaceutical Sciences, and 0421 and A2022IQB001-06 in the Institute for Quantitative Biosciences). All procedures were followed in accordance with The University of Tokyo guidelines for the care and use of laboratory animals and ARRIVE guidelines.

### Mouse maintenance and preparation of pregnant mice

JCL:ICR (CLEA Japan), Slc:ICR (Japan SLC), and C57BL/6J mice were housed in a temperature- and humidity-controlled environment (23 ± 3 °C and 50 ± 15%, respectively) under a 12-h light/dark cycle. Animals were housed in sterile cages (Innocage, Innovive) containing bedding chips (PALSOFT, Oriental Yeast), with two to six mice per cage, and provided irradiated food (CE-2, CLEA Japan) and filtered water ad libitum. For in utero electroporation experiments, mating was limited to a 2-h period, and the presence of a vaginal plug was used to confirm the onset of gestation (E0.0).

### Plasmid constructs

pAAV-DIO-GFP (pAAV-Ef1a-DIO-EGFP-WPRE-pA) and pNeuroD1-IRES-GFP were kindly provided by Frank Polleux (Jossin and Cooper, 2011). For the construction of pNeuroD1-ERT2CreERT2, ERT2CreERT2 was digested from pCAG-ERT2CreERT2 (Addgene, #13777) and cloned into pNeuroD1-IRES-GPF digested with the same restriction enzymes. pCAG-mCherry was constructed in the previous report (Kishi et al., 2023).

### In utero electroporation

E12.0 mice were administered a sodium pentobarbital–based anesthetic, and the uterine horn was exposed. The embryo was pinched between a flexible fiber-optic cable and a finger, and plasmid DNA (pNeuroD1-ERT2CreERT2, 0.5 μg/μL; pAAV-DIO-GFP, 1 μg/μL; pNeuroD1-IRES-GFP, 0.5 μg/μL; pCAG-mCherry, 1 μg/μL) with 0.01% FastGreen was injected into the lateral ventricle. The uterus was held between tweezer-type electrodes (CUY650P3, Nepa Gene) and electroporated with CUY21EDIT or NEPA21 system (Nepa Gene), according to the program: four pulses of 30 V with a 50-ms pulse width and 950-ms pulse interval. After injection and electroporation, the uterine horn was returned to its original location. The mice recovered on a heating plate maintained at 38 °C until they regained consciousness following anesthesia.

### Immunohistochemistry

Immunofluorescence staining was performed as described previously (Eto et al., 2020). E13.0 embryos were directly fixed with 4% paraformaldehyde (PFA) in PBS at 4 °C for 2–3 h. E14.0 and E16.0 embryos were perfused with 4% PFA in PBS, and isolated brains were postfixed with 4% PFA in PBS at 4 °C for 2–3 h. After equilibration with 30% (w/v) sucrose in PBS, the fixed brains were embedded in an OCT compound (Tissue TEK) and frozen. Coronal cryosections (∼15 μm) on slide glasses were exposed to TBS with 0.1% Triton-X-100 (TBS-T) and 3% BSA (Blocking buffer) and incubated with anti-GFP antibody (Nakalai, GF090R, 1:1,000 dilution) and anti-DsRed antibody (MBL, PM005, dilution 1:000) for mCherry in blocking buffer at 4 °C overnight. After washing with TBS-T, the slide glasses were incubated with Alexa Fluor–conjugated secondary antibodies in a blocking buffer and mounted in Mowiol (Calbiochem). All fluorescent images were obtained with laser confocal microscopy (Leica TCS-SP5) and analyzed using the ImageJ software (NIH).

### Isolation of NPCs and GFP-labeled neurons by FACS

Isolation of NPCs and neurons was performed using a previously established protocol (Kishi and Gotoh, 2021). The neocortices of non-electroporated E12.0 or E13.0, E14,0, and E16.0 after electroporation were dissected and subjected to enzymatic digestion using Neuron Dissociation Solution (FUJIFILM Wako Chemicals). For E12.0 NPCs, dissociated cells were stained with allophycocyanin (APC)-conjugated anti-CD133 antibody (BioLegend, 141208). E12.0 NPCs as CD133-high cells or E13.0, E14.0, and E16.0 neurons as GFP-positive cells were isolated using the FACSAria instrument (Becton Dickinson).

### RT-qPCR

RT-qPCR was performed as described previously (Kishi and Gotoh, 2021). Total RNA was extracted from isolated cells by FACS with RNAiso Plus (Takara Bio) and subjected to reverse transcription with ReverTra Ace qPCR RT Master Mix with gDNA Remover (Toyobo). The concentrations of target cDNA were determined with THUNDERBIRD SYBR qPCR Mix (Toyobo) using LightCycler 480 or LC96 instruments (Roche). The expression levels were calculated by absolute quantification and normalized with *the Actb* housekeeping gene. Primer sequences are as follows: *Actb*, AATAGTCATTCCAAGTATCCATGAAA, and GCGACCATCCTCCTCTTAG; *Nestin*, TGAAGCACTGGGAAGAGTA, and TAACTCATCTGCCTCACTGTC; *NeuroD1*, TACGACATGAACGGCTGCTA, and TCTCCACCTTGCTCACTTT; *Tubb3*, ACACAGACGAGACCTACT, and GCAGACACAAGGTGGTT; *Gabrb2*, ATGCCATCAATTCTGATTACCA, and TAATTCCTAATGCAACCCGTG.

### RNA-seq

Libraries for RNA-seq analysis were constructed from total RNA isolated from two independent experiments at each stage as for RT-qPCR analysis. A TruSeq RNA sample preparation kit (Illumina) was used for template preparation, which was followed by sequencing using the HiSeq platform (Illumina) to obtain 100-base paired-end reads. More than 20 million fragments were analyzed. For layer-specific gene expression analysis, we obtained raw sequence files from the GEO database (GSE27243; GSM673634 for layers 1–3 and GSM673639 for layer 6) (Belgard et al., 2011). Adaptor sequences, pNeuroD1 plasmid–containing sequences, low-quality sequences (MAPQ < 30), and poly-A sequences were removed with Trimmomatic, bowtie, Fastx-toolkit, and PRINSEQ software, respectively (Bolger et al., 2014; Langmead et al., 2009; Schmieder and Edwards, 2011). Reads were mapped to the reference mouse genome (mm10) with HISAT2 (Kim et al., 2019). Mapped reads to ribosomal RNA genes and sex chromosomes were removed with samtools and bedtools (Li et al., 2009; Quinlan and Hall, 2010), since pooled cells from both male and female embryos were used in this study. Reads mapped on protein-coding genes based on GENCODE vM20 mouse genome reference (Frankish et al., 2018) were counted with featureCounts and normalized with iDEGS/edgeR (Liao et al., 2014; Robinson et al., 2010). To compare the expression levels between different genes, the values were normalized by the gene length, and this value was called “expression value” in this study. Differentially expressed genes were determined as genes with FDR < 0.05 calculated with iDEGES/edgeR. GO analysis was performed using DAVID (Huang et al., 2009a, 2009b) (Huang 2009 x 2). GSEA analysis was performed using software provided by Broad Institute (Mootha et al., 2003; Subramanian et al., 2005).

### DNase-seq

Isolated cells from two independent experiments by FACS were resuspended with nuclear buffer A (85mM KCl, 5.5% Sucrose, 10mM Tris-HCl (pH 7.5), 0.5mM Spermidine, 0.2mM EDTA) and gently mixed with the same volume of nuclear buffer B (Nuclear buffer A with 0.1% NP-40). After centrifuging at 600 g for 5 min at 4 °C and removing supernatant, pelleted nuclei were resuspended with nuclear buffer R (85 mM KCl, 5.5% sucrose, 10 mM Tris-HCl (pH 7.5), 3 mM MgCl2, 1.5 mM CaCl2), added with DNaseI (Takara Bio) at a final concentration of 2 U/mL, and incubated for 30 min at 37 °C. After quenching the reaction by adding lysis buffer (1% SDS, 50mM Tris-HCl (pH 8.0), 10mM EDTA), digested DNA was purified by phenol-chloroform–isoamyl alcohol and subjected to agarose gel electrophoresis. DNA less than one kbp was purified by FastGene Gel/PCR Extraction kit (NIPPON Genetics).

Libraries for DNase-seq analysis were constructed by a TruSeq ChIP sample preparation kit (Illumina), which was used for template preparation, followed by sequencing using the HiSeq platform (Illumina) to obtain 36 base-paired-end reads. More than 20 million fragments were analyzed. pNeuroD1 and pAAV-DIO-EGFP plasmid– containing sequences were removed with bowtie software. Reads were mapped to the reference mouse genome (mm10) with bowtie, allowing a maximum of two mismatches. Only the uniquely aligned reads were used for further analysis. Sequenced reads mapped to known ENCODE blacklist regions (Amemiya et al., 2019; Consortium et al., 2012) and sex chromosomes were removed by bedtools and samtools, respectively. The peaks were determined by F-seq as a region with *p* < 10^−6^, and the peaks detected in both two replicates were determined as DHSs. Opening or closing regions were determined as DHSs newly appeared or disappeared during differentiation, respectively.

To determine the genome features, the protein-coding genes annotated in GENCODE vM20 were used. Promoters were determined to be 1 kb upstream and 0.5 kb downstream of the transcription start sites, and exons and introns did not include promoters. Downstream was determined as 1 kb downstream of the transcription terminate site without promoters, exons, and introns. Distal regions were determined as the regions that had not been characterized. ChIP-Atlas was utilized to find the transcription factors that especially bind to DHSs determined in E16.0 neurons (Oki et al., 2018).

### Microarray

The cortex from E15 C57BL/6J mice was isolated, dissociated with neuron dissociation solutions (FUJIFILM Wako Chemicals), and plated at a density of approximately 1.2 cells/cm^2^ on poly-D-lysine–coated dishes in neurobasal medium (Gibco) supplemented with 1% Gultamax (Gibco) and 2% B27 (Gibco). After 6 days, the culture was treated with 50 ng/mL BDNF (Sigma-Aldrich) for 1 h. The RNA isolated for RT-qPCR was subjected to analysis using SurePrint G3 Mouse GE 8x60K Microarray (Agilent), according to the manufacturer’s instructions (Agilent SureScan). The data was normalized by percentile shift (75%) method with GeneSpring software.

### ChIP-seq

ChIP was performed as described previously (Eto et al., 2020). Isolated NPCs from two independent experiments by FACS from E12.0 embryos were fixed with 1% formaldehyde and stored at −80 °C until the experiment. The cells were thawed, suspended in RIPA buffer for sonication (10 mM Tris HCl at pH 8.0, 1 mM EDTA, 140 mM NaCl, 1% Triton X 100, 0.1% SDS, and 0.1% sodium deoxycholate), and subjected to ultrasonic treatment using Picoruptor (15 cycles of 30 s ON and 30 s OFF) (Diagenode). The cell lysates were then diluted with RIPA buffer for immunoprecipitation (50 mM Tris-HCl at pH 8.0, 150 mM NaCl, 2 mM EDTA, 1% Nonidet P 40, 0.1% SDS, and 0.5% sodium deoxycholate), incubated for 1 h at 4 °C with protein A/G magnetic beads (Pierce) to clear non-specific binding, and then incubated overnight at 4 °C with protein A/G magnetic beads that had previously been incubated overnight at 4 °C with 2 μL of antibodies to H3K27me3 (MBL, MABI0323) or H3K4me3 (MBL, MABI0304). The beads were isolated and washed three times with wash buffer (2 mM EDTA, 150 mM NaCl, 0.1% SDS, 1% Triton X 100, and 20 mM Tris HCl at pH 8.0) and then once with wash buffer containing 500 mM NaCl. Immune complexes were eluted from the beads for 15 min at 65 °C in Direct Elution buffer (10 mM Tris HCl at pH 8.0, 5 mM EDTA, 300 mM NaCl, and 0.5% SDS), and they were then subjected to digestion with proteinase K (Nacalai Tesque) for > 6 h at 37 °C, and decrosslinked by incubation at 65 °C for > 6 h. Immunoprecipitated DNA was purified with phenol–chloroform–isoamyl alcohol.

Libraries for ChIP-seq analysis were constructed with a TruSeq ChIP sample preparation kit (Illumina), which was used for template preparation, followed by sequencing using the HiSeq platform (Illumina) to obtain 36 base-paired-end reads. More than 20 million fragments were analyzed. Processing, mapping, and peak calling were the same as for DNase-seq. The regions that were determined as peaks in both two replicates were determined as the regions with histone modification. Genes were classified into five clusters with histone modification profiles by k-means clustering by ngsplot (Shen et al., 2014), and genes with both H3K4me3 and H3K27me3 modifications were determined as bivalent genes in E12.0 NPC.

### Motif enrichment analysis

Motif enrichment analysis was performed by HOMER against background sequences randomly selected from the whole genome with similar GC % (except for repeat sequences) (Heinz et al., 2010).

### Statistical analysis

Data were compared using an analysis of variance (ANOVA), followed by Tukey’s multiple comparison test or Friedman’s test, followed by Dunn’s multiple comparison test. A *p*-value of <0.05 was considered statistically significant.

## Supporting information

Sapplemental Data 1

Sapplemental Data 2

Sapplemental Data 3

Sapplemental Data 4

## Acknowledgments

We would like to thank F. Polleux (Columbia University) for providing the plasmid pNeuroD1-IRES-GFP; N. Shinoda, Y. Yamaguchi, and M. Miura (The University of Tokyo) for helping with microarray analysis; R. Nakato, K. Kadota, T. Horiuchi (The University of Tokyo), and S. Oki (Kumamoto University) for advising regarding sequence analysis; the Human Genome Center (The University of Tokyo) for providing the super-computing resource; the One-Stop Sharing Facility Center for Future Drug Discoveries (The University of Tokyo) for providing FACS; R. Nagayoshi, Y. Kakeya, and M. Saeki (The University of Tokyo) for technical assistance; and members of the Gotoh and Kishi laboratories for helpful discussion. This research was supported by AMED-CREST (JP23gm1310004 to Y.G.), AMED-PRIME (JP22gm6110021 to Y.K.), MEXT/JSPS KAKENHI (JP22H00431 to Y.G.; 16H06279, JP22H04687, 23H04214, and 24H01227 to Y.K.), the Takeda Science Foundation, the Uehara Memorial Foundation, the Asahi Glass Foundation, the Chugai Foundation for Innovative Drug Discovery Science, the Astellas Foundation for Research on Metabolic Disorders, the Naito Foundation, and the SECOM Science and Technology Foundation.

## Author Contributions

Conceptualization, S.S., Y.K., and Y.G.; Data Curation, S.S., Y.K., and Y.S.; Formal Analysis; S.S. and Y.K.; Funding Acquisition, S.S., Y.K., and Y.G.; Investigation, S.S., Y.K., Y.M., K.K., and Y.S.; Methodology, Y.K. and Y.M.; Project administration, Y.K. and Y.G.; Supervision, Y.K. and Y.G.; Visualization, S.S. and Y.K; Writing – Original Draft Preparation; S.S. and Y.K.; Writing – Review & Editing, S.S., Y.K., and Y.G.

## Data availability

The sequence and microarray data have been deposited in the DNA Data Bank of Japan (DDBJ) Sequence Read Archive under the following accession codes: DRA015915 (RNA-seq), DRA015284, and DRA018693 (DNase-seq), and DRA018694 (H3K4me3 and H3K27me3 ChIP-seq). The processed data of the sequences and microarray data have also been deposited in the DDBJ Gene Expression Archive under the following accession codes: E-GEAD-803 (microarray of BDNF stimulation), E-GEAD-859 (peak files of DHSs), and E-GEAD-860 (bigwig files of ChIP-seq). We also used published datasets: GSE27243 (layer-specific RNA-seq in adult brain) (Belgard et al., 2011).

## Figure legends

**Supplementary Figure 1.**
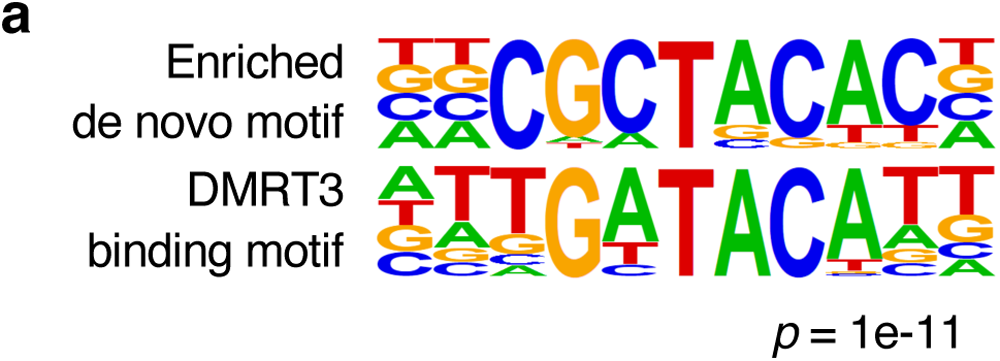
Enrichment of DNA sequence recognized by Dnmt3 in E12.0–E16.0 opening and bivalent regions. **(a)** Motif analysis for E12.0–E16.0 opening and bivalent regions by HOMER software. Enriched motif (upper panel) and Dmrt3-binding motif (lower panel) were shown.

**Supplementary Data 1. Transcriptome analysis, chromatin accessibility, and epigenetic state during differentiation of deep-layer neurons, related to Figures 2–6**.

The expression levels, the presence of DHS in promoter, upper- or deep-layer specific expression, and the state of H3K4me3 and H3K27me3 of all genes are shown.

**Supplementary Data 2. The expression changes of E12 to E16 opening and layer 6-specific genes during embryonic to adult stage, related to** Figure 5.

**Supplementary Data 3. Microarray analysis of primary neuronal culture with or without BDNF treatment, related to** Figure 5.

**Supplementary Data 4. ChIP-atlas analysis of E16 DHSs, related to** Figure 6.

